# Biochemical and physiological investigations on adenosine 5’ monophosphate deaminase from Plasmodium spp.

**DOI:** 10.1101/447789

**Authors:** Lakshmeesha Kempaiah Nagappa, Hemalatha Balaram

**Affiliations:** Molecular Biology and Genetics Unit, Jawaharlal Nehru Centre for Advanced Scientific Research (JNCASR), Bengaluru, Karnataka, INDIA.

**Keywords:** *Plasmodium*, AMP deaminase, functional complementation in yeast, asexual stage viability, adenylate energy charge

## Abstract

HGXPRT - Hypoxanthine-guanine-xanthine phosphoribosyltransferase, ADSS - Adenylosuccinate synthetase, ASL - Adenylosuccinate lyase, GMPS - Guonosine monophosphate synthetase, IMPDH - Inosine monophosphate dehydrogenase, ISN1 - Inosine monophosphate specific nucleotidase, PNP - Purine nucleoside phosphorylase

**Summary:** Interplay between ATP generating and utilizing pathways in a cell is responsible for maintaining cellular ATP/energy homeostasis that is reflected by Adenylate Energy Charge (AEC) ratio. Adenylate kinase (AK), that catalyzes inter-conversion of ADP, ATP and AMP, plays a major role in maintaining AEC, and is regulated by cellular AMP levels. Hence, the enzymes AMP deaminase (AMPD) and nucleotidases, which catabolize AMP, indirectly regulate AK activity and in-turn affect AEC. Here, we present the first report on AMPD from *Plasmodium*, the causative agent of malaria. The recombinant enzyme expressed in *Saccharomyces cerevisiae* was studied using functional complementation assay and residues vital for enzyme activity have been identified. Similarities and differences between *Plasmodium falciparum* AMPD (PfAMPD) and its homologs from yeast, *Arabidopsis* and humans are also discussed. The AMPD gene was deleted in the murine malaria parasite *P. berghei* and was found to be non-essential for intra-erythrocytic growth of the knockout parasites. However, when episomal expression was attempted, viable parasites were not obtained, suggesting that perturbing AMP homeostasis by over-expressing AMPD might be lethal. As AMPD is known to be allosterically modulated by ATP, GTP and phosphate, allosteric activators of PfAMPD could be developed as anti-parasitic agents.

## 1 Introduction

Malaria caused by the parasitic protozoan *Plasmodium* is one of the deadliest diseases causing 445,000 deaths per annum worldwide (WHO, 2017). Although the PlasmoDB database is replete with information on the genome and proteome of the organism, most proteins are uncharacterized or have putative status (Aurrecoechea et al., 2008; Bahl et al., 2002, 2003). This inadequacy of proper understanding of the biochemistry of the organism has been a major impediment to the efforts put forth towards mitigation of this menacing disease. With genetic manipulation being highly challenging in *P. falciparum*, understanding the physiological significance of these uncharacterized/putative proteins becomes a non-trivial task (de Koning-Ward et al., 2000). As the parasite is completely dependent on salvage of precursors (predominantly hypoxanthine) for its purine nucleotide bio-synthesis (Downie et al., 2008), the enzymes involved in this pathway have been sought after as potential drug targets. Although many enzymes of the purine nucleotide metabolism in *Plasmodium* have been studied in great detail (Hypoxanthine-guanine-xanthine phosphoribosyltransferase (HGXPRT), adenylosuccinate synthetase (ADSS), adenylosuccinatelyase (ASL), purine nucleoside phosphorylase) (Jayalakshmi et al., 2002; Eaazhisai et al., 2004; Bulusu et al., 2009; Mehrotra et al., 2010; Roy et al., 2015a,b; Madrid et al., 2008), a few more remain uncharacterized (AMPD, IMP dehydrogenase, IMP-specific nucleotidase and other purine nucleotidases). Through this study, we add further knowledge to the existing understanding of this protozoan parasite by examining one more enzyme involved in purine nucleotide metabolism, AMP deaminase (EC. 3.5.4.6), a previously uncharacterized protein, which has been implicated in physiological disorders such as myoadenylate deaminase deficiency (a muscular disorder) and pontocerebellar hypoplasia (a neurological disorder) (Fishbein et al., 1978; Shumate et al., 1979; Morisaki et al., 1992; Hellsten et al., 1999; Akizu et al., 2013). AMP deaminase has also been considered as a herbicide target and inhibition of its isoform in humans has been known to provide relief during myocardial infarction (Sabina et al., 2007; Bazin et al., 2003).

AMPD belongs to amidohydrolase superfamily of enzymes characterized by a conserved (*α/β*)_8_- barrel core and a mononuclear metal centre (Seibert and Raushel, 2005). Salient features of AMPD in particular include, exclusive eukaryotic occurrence, presence of a divalent zinc at the metal centre and a highly divergent N-terminus (Sabina and Mahnke-Zizelman, 2000; Marotta et al., 2009). Although detailed biochemical characterization of *S. cerevisiae* AMPD (ScAMPD) (Yoshino et al., 1979; Murakami, 1979; Murakami et al., 1980; Meyer et al., 1989; Merkler et al., 1989; Merkler and Schramm, 1990, 1993), and other AMPDs (from rabbit skeletal muscle, gold fish, sea scorpion etc.) have been carried out earlier (Marotta et al., 2009; Waarde and Kesbeke, 1981; Lushchak et al., 1998; Mangani et al., 2003; Martini et al., 2007), the only crystal structure available in the PDB database for this enzyme is from *Arabidopsis thaliana*, solved in 2006 by the group of Richard L Sabina (Han et al., 2006). Recently, Akizu *et al.,* identified mutations in AMPD2 in families where individuals were found to be suffering from a neurodegenerative brainstem disorder termed ‘pontocerebellar hypoplasia’ (Akizu et al., 2013). In addition to this, a new mechanism of action for the anti-diabetic drug metformin has been proposed by Ouyang *et al.,* where it was shown that AMPD inhibition in L6 rat skeletal muscle cells by metformin, led to an increase in AMP levels which in-turn activated AMP activated protein kinase (AMPK) (Ouyang et al., 2011). Also, AMPD and AMPK have been reported to counter-regulate each other and are implicated in hepatic steatosis (Lanaspa et al., 2012).

AMPD can modulate AEC in a dual manner by being part of purine nucleotide cycle as well as by independently regulating the equilibrium of the adenylate kinase reaction. Adenylate kinase catalyses a reversible reaction; where two ADP molecules are converted to one ATP and one AMP (2ADP *↔* ATP + AMP) (Atkinson, 1968; Chapman and Atkinson, 1973). This reaction maintains the three way equilibrium between ATP, ADP and AMP levels in a cell. The ATP generated will be utilized by energy consuming processes, while AMP gets deaminated to IMP by AMPD (Scheme 1). In conditions where AMPD is not operating (either due to a mutation in case of the genetic disorder or a knock-out where the gene has been deleted) AMP accumulation takes place and this adversely affects the adenylate kinase reaction by shifting the reaction equilibrium towards ATP depletion. This leads to several consequences of which, most prominent in the literature is fatigue and muscle weakness reported in subjects suffering from myoadenylate deaminase deficiency, an autosomal recessive genetic disorder (Hellsten et al., 1999; Isackson et al., 2005). In the absence of AMPD activity insufficient production of ATP and accumulation of AMP synergistically bring down the AEC of the cell. Further, in patients suffering from pontocerebellar hypoplasia, lack of AMPD activity results in accumulation of AMP which leads to inhibition of *de novo* purine biosynthesis pathway which eventually manifests in impaired protein production due to insufficient GTP pools. Also, AMPD2 and AMPD3 double knockout in mice has been found to be post-nataly lethal, while mutation in AMPD gene (FAC1) was found to be embryonically lethal in *Arabidopsis*, making it a good herbicide target (Akizu et al., 2013; Sabina et al., 2007; Xu et al., 2005).

**Scheme 1:**
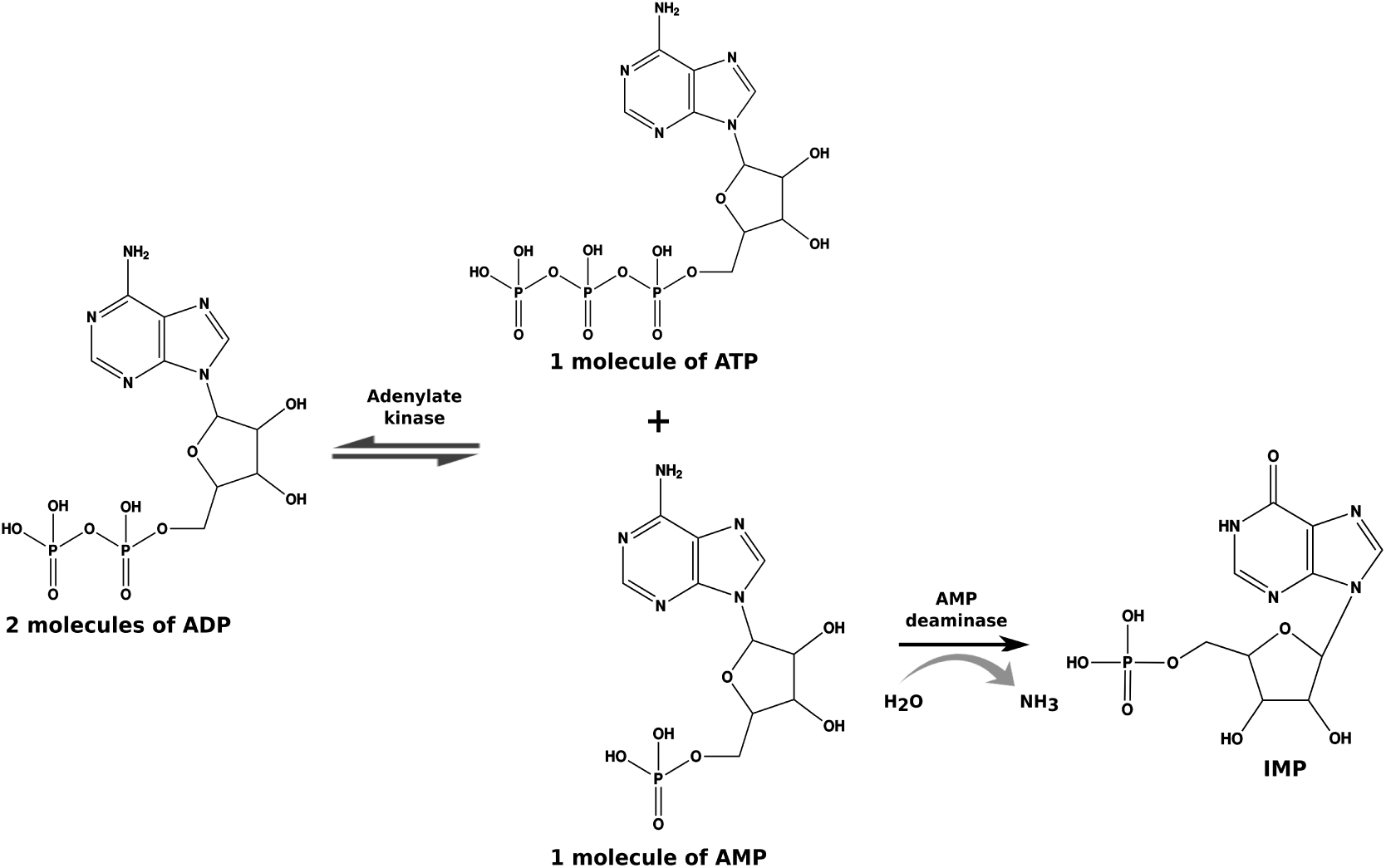
Adenylate kinase and AMP deaminase reaction. Adenylate kinase catalyses the formation of ATP and AMP using two ADP molecules. ATP fuels energy utilizing cellular processes where as AMP can be deaminated to IMP by AMP deaminase with concomitant release of ammonia.

Studies have documented increase in AMPD activity in the soil-living amoeba *Dictiostelium discoidium* in response to nutrient starvation or treatment with hadacidin (inhibitor of ADSS) (Jahngen and Rossomando, 1986). It was also found that loss of AMPD function resulted in increased AMP secretion, impaired ratio of prestalk to prespore cells and reduced sporulation efficiency (Chae et al., 2002). In the context of malaria, a gain-of-function mutation in erythrocytic AMPD was found to increase RBC turnover and confer resistance in mice against *P. chabaudi* infection (Hortle et al., 2016). Also, an indirect evidence for the presence of AMPD activity in *Plasmodium* was provided by Cassera *et al.,* where it was shown that erythrocytic AMP can be taken up by the parasite and converted to IMP by the action of AMPD (Cassera et al., 2008). But, a detailed biochemical and physiological characterization of this enzyme from the parasite has remained hitherto unavailable. In the current report we provide a comprehensive biochemical description of the *P. falciparum* enzyme, making judicious use of the information from the crystal structure of *Arabidopsis thaliana* AMPD (AtAMPD) and forward genetics studies conducted on human AMPD2 as well as the *Arabidopsis* enzyme (Akizu et al., 2013; Han et al., 2006; Xu et al., 2005; Hortle et al., 2016). We have used a functional complementation assay to identify and validate essentiality of key conserved residues required for this 693 amino acid protein to remain functional. Our results throw light upon some of the key features of *P. falciparum* AMPD (PfAMPD) in comparison with human, *Arabidopsis* and yeast counterparts. Further, the *in vivo* genetic manipulation carried out in *P. berghei* (Pb), involving genetic ablation and GFP tagging provides evidence for the dispensable nature of this cytosolic enzyme during the intra-erythrocytic stages, while expression of wildtype GFP-tagged *P. berghei* AMPD (PbAMPD) protein through an episomal construct was found to be toxic.

## 2 Materials

Chemicals and reagents were procured from Sigma Aldrich, USA; Fischer Scientific; SRL, India. Media components were purchased from Himedia, Mumbai, India. Primers were custom synthesized from Sigma Aldrich, Bengaluru, India. Restriction enzymes, reverse transcriptase, Phusion DNA polymerase, Taq DNA polymerase and T4 DNA ligase were obtained from New England Biolabs, USA and used according to the instructions of the manufacturer. Trizol� was from Invitrogen, USA. Antibiotics ampicillin, tetracycline and chloramphenicol were bought from USB chemicals. G418 was bought from Sigma Aldrich, USA and Zeocin from Invitrogen, USA. Plasmid isolation, PCR clean-up and gel extraction kits were from Qiagen. *E. coli* strain XL-1 blue, and expression strain Rosetta (DE3) pLysS and plasmid pETduet-1 were from Novagen. pJAZZ library clone (PbG02_B-44h05); plasmids, pSC101BAD, R6K Zeo/pheS and *E. coli* pir strains were procured from Plasmo-Gem, Sanger Institute, UK. *E. coli* TSA cells were obtained from Lucigen. Plasmid pBCEN5 was a gift from Dr. Shiroh Iwanaga, Japan. Wild type *(BY4742; MATα; ura3*∆*0; leu2*∆*0; his3*∆*1; lys2*∆*0)*, ∆*amd1 (BY4742; MATα; ura3*∆*0; leu2*∆*0; his3*∆*1; lys2*∆*0; YML035c::kanMX4),* ∆*ade1 (BY4741; MATa; ura3*∆*0; leu2*∆*0; his3*∆*1; met15*∆*0; YAR015W::kanMX4)* and ∆*ade2 (BY4741; MATa; ura3*∆*0; leu2*∆*0; his3*∆*1; met15*∆*0; YOR128C::kanMX4) S. cerevisiae* strains were from EUROSCARF, Germany. Plasmid pYES2C/T was from Invitrogen, USA. Codon harmonised PfAMPD gene was synthesized and procured from Shinegene, China. *P. falciparum* 3D7 and *P. berghei* ANKA strain were procured from MR4. RPMI-1640 and Albumax-I were from Sigma Aldrich, USA and Gibco, respectively. Amaxa 4D nucleofector and P5 Nucleofection kit were from Lonza, Germany. Infusion assembly kit was from TaKaRa Bio, USA. Random primer kit was from BRIT, India. Male/female BALB/c mice aged 6-8 weeks were used for cultivation and transfection of *P. berghei*. Gene sequencing by Sanger sequencing method was carried out in the in-house central instrumentation facility, JNCASR, Bangalore. Sequences of oligonucleotides used are provided in the section ‘Supplementary information’ (Table S1).

## 3 Experimental Procedures

### 3.1 RNA isolation, cDNA synthesis and RT-PCR

Saponin released *P. falciparum* cell pellet from 40 mL of *in vitro* culture was treated with 1 mL Trizol� reagent at room temperature for 15 minutes and then centrifuged for 30 minutes at 18000 × g to remove the cell debris. The supernatant was transferred to a fresh tube and 200 *μ*L of chloroform was added and kept at room temperature for 15 minutes. This was centrifuged at 18000 × g for 30 minutes at 4 °C for phase separation. The top layer was aspirated to a fresh tube and 400 *μ*L of isopropanol was added and kept at −20 °C for 4 hours. This was centrifuged at 18000 × g for 30 minutes at 4 °C to obtain a RNA pellet. This pellet was washed once with 75 % ethanol made in diethylpyrocarbonate (DEPC) treated water and allowed to air dry. The pellet was resuspended in 20 *μ*L of DEPC treated water and stored at −80 °C. For RNA isolation from *P. berghei*, blood was harvested from mice at 10 – 15 % parasitemia and collected in heparin (200 units mL*^−^*^1^). The cells were lysed by resuspending them in 10 volumes of erythrocyte lysis buffer (155 mM NH_4_Cl, 12 mM NaHCO_3_ and 0.1 mM EDTA) and kept at 4 °C for 5 minutes. The cells were harvested by centrifugation at 500 × g for 10 minutes, the supernatant was discarded and the pellet was used for RNA isolation. The steps used for RNA isolation were identical to that described for *P. falciparum*. The cDNA was synthesized using gene specific reverse primer (P1 for *P. falciparum* and P73 for *P. berghei*) and Moloney Murine Leukemia Virus (MMLV) reverse transcriptase according to manufacturer's protocol. This cDNA was used as template for PCR amplification of AMPD gene.

### 3.2 Generation of plasmid constructs

All clones were generated in XL-1 blue strain of *E. coli* cells (Table S2 lists the plasmid constructs used in the study). Using cDNA as template PfAMPD gene was PCR amplified using primers P2 and P3. The PCR product was digested with EcoRI and HindIII and cloned into double digested pETDuet-1 vector. Similarly, the truncated versions of PfAMPD gene were PCR amplified using the full length clone as template and primer pairs P4, P5 for ∆59; P6, P7 for ∆94 and P8, P9 for ∆178. All PCR products were digested using EcoRI and HindIII enzymes and cloned into double digested pETDuet-1 vector. Using genomic DNA from yeast as template, yeast AMPD gene was PCR amplified using primers P10 and P11. The PCR product was digested with SacI and XhoI enzymes and cloned into double digested pYES2/CT plasmid. Using full length clone of PfAMPD (pETDuet-1_PfAMPD) as template and primers P12, P13 PfAMPD gene was PCR amplified, digested with KpnI and XbaI enzymes followed by cloning into double digested pYES2/CT plasmid. Harmonised PfAMPD gene segment was released from pMD19_hPfAMPD vector using the enzymes KpnI and XhoI and subcloned into pYES2/CT plasmid which was cut using the same restriction sites. eGFP was amplified using primers P14, P15 and pRS306 as template, double digested using enzymes BamHI and XhoI and ligated into double digested pYES2/CT vector. Cloning of GFP tagged yeast and PfAMPD was done by ligating three DNA segments i.e., vector, gene fragment and GFP fragment. ScAMPD was amplified using primers P16, P17 followed by digestion with SacI and KasI enzymes. eGFP was amplified by using primers P18, P19 followed by digestion with KasI and XhoI enzymes. These two fragments were ligated with SacI, XhoI digested pYES2/CT plasmid. Similarly PfAMPD was amplified using primers P20, P21 and pMD19_hAMPD as template, followed by digestion with KpnI and XbaI enzymes. eGFP was amplified by using primers P22, P23 followed by digestion with XbaI and XhoI enzymes. These two fragments were ligated with KpnI, XhoI digested pYES2/CT plasmid. PCR product for pYES2/CT_(His)_6__hPfAMPD_GFP construct was generated using primers P56, P55 and pYES2/CT_hPfAMPD_GFP (pYES2/CT clone containing harmonised PfAMPD gene with C-terminal GFP tag) as template. The PCR product was digested with enzymes KpnI and XhoI and cloned into digested pYES2/CT plasmid. PCR product for pYES2/CT_hPfAMPD_GFP_(His)_6_ construct was generated using primers P54, P57 and pYES2/CT_hPfAMPD_GFP as template. The PCR product was digested with enzymes KpnI and XhoI and cloned into digested pYES2/CT plasmid.

### 3.3 Bacterial transformation and expression

Competent *E. coli* cells were prepared by following the method described by Chung *et al.,* Trasformation was carried out as described in ‘Molecular cloning’ by Sambrook and Russel (Sambrook and Russell, 2006). For expression, single colony was inocluated into 5 mL LB media with ampicillin (100 *μ*g mL*^−^*^1^) and chloramphenicol (34 *μ*g mL*^−^*^1^) and grown overnight at 37 °C, 180 rpm. The overnight culture was added to a secondary culture (TB) at 1 % inoculum in the presence of the antibiotics ampicillin and chloramphenicol and incubated at 37 °C, 180 rpm till the OD_600_ reached 0.6. The culture was then induced using IPTG at a concentration of 1 mM and further incubated at 37 °C, 180 rpm for four more hours. 1 mL culture was lysed by resuspending in 1 × loading dye and separated on SDS-PAGE followed by Western blot. Anti - (His)_6_ antibody from mouse was used to probe the blot followed by anti - mouse antibody from goat conjugated to horse radish peroxidase (HRP) as secondary antibody. The blot was developed using amino ethyl carbazole (AEC) and hydrogen peroxide as substrates.

### 3.4 Yeast transformation and expression

*Saccharomyces cerevisiae* trasformation was performed according to the protocol described by Gietz *et al.,* (Gietz and Woods, 2002). For expression, single colony was inocluated into 5 mL SD-Ura (synthetically defined minus uracil) medium containing glucose (2%) as carbon source and incubated at 30 °C, 180 rpm for 15-20 hours. The overnight culture was harvested by centrifugation, washed once with sterile water and resuspended in 5 mL of SD-Ura medium containing galactose (2 %) as carbon source and incubated at 30 °C, 180 rpm. 1 mL aliquots were taken at different time points and TCA was added to the cells (final concentration 12 %) and stored at −80 °C overnight for precipitation. The aliquots were thawed and the cells were harvested by centrifugation at 12000 × g. The supernantant was discarded and the pellet was washed in 250 *μ*L 80 % cold acetone twice. After the second wash, acetone was removed and the pellet was air dried, resuspended in 1 % SDS/0.1 N NaOH solution. The lysates were boiled in the presence of 1 × loading dye and separated on SDS-PAGE followed by Western blotting. Mouse anti - (His)_6_ antibody or anti - GFP antibody was used to probe the blot followed by anti - mouse antibody from goat conjugated to (HRP) as secondary antibody. The blot was developed using AEC and hydrogen peroxide as substrates.

### 3.5 Serial dilution and spotting of *S. cerevisiae* cells

Single colonies were inoculated in to 5 mL SD-Ura medium containing glucose as carbon source and grown overnight at 30 °C, 180 rpm. The cells were harvested by centrifugation at 5000 × g and washed once with sterile water. OD_600_ was determined and serial dilutions were made starting from 1 OD mL*^−^*^1^ to 10*^−^*^3^ OD mL*^−^*^1^. 2 *μ*L from each dilution was spotted on SD-Ura plate containing galactose as carbon source with or without 50 *μ*g mL*^−^*^1^ S - adenosyl methionine (SAM). The plates were allowed to air dry and incubated at 30 °C (37 °C for temperature sensitivity studies) for 60-72 hours. The plates were observed for cell growth and photographically documented.

### 3.6 Site-directed mutagenesis and PfAMPD truncations

Using pYES2/CT_hPfAMPD_GFP (pYES2/CT clone containing harmonised PfAMPD gene with C-terminal GFP tag) plasmid as template site-directed mutagenesis was performed by following overlap extension PCR and AQUA cloning protocol (Beyer et al., 2015). Partial amplicons for each mutant were obtained by PCR using Gal promoter-forward primer (P54)/ mutagenic-reverse primer and Cyc terminator-reverse primer (P55)/ mutagenic-forward primer pairs (Table S1). The fragments were gel purified and mixed with double digested (KpnI and XhoI) and gel purified pYES2/CT vector in molar ratio 1:3:3:: vector:insert1:insert2. This mix was used to transform competent XL1-Blue *E. coli* cells and plated on LB plates containing 100 *μ*g mL*^−^*^1^ ampicillin. The clones were confirmed by PCR, restriction digestion and sequencing. These constructs were used to perform functional complementation in ∆*amd1* yeast by serial dilution and spotting assay. PfAMPD lacking 60 and 180 residues from N-terminal end, hPfAMPD*_*∆*N60* and hPfAMPD*_*∆*N180* were PCR amplified using primer pairs P24, P55 and P25, P55 (Table S1); respectively. The PCR product was double digested (KpnI and XhoI) and cloned into digested pYES2/CT plasmid.

### 3.7 Generation of *P. berghei* transfection vectors

The library clone for *P. berghei* AMPD (PbG02_B-44h05) was obtained from PlasmoGem. The procedures for knockout construct generation were described earlier (Godiska et al., 2009; Pfander et al., 2011). The intermediate vector and final knockout construct were confirmed by PCR and restriction digestion. The final knockout construct was subjected to NotI digestion, purified and used for transfection. For episomal expression of GFP-tagged PbAMPD, *P. berghei* AMPD gene was amplified by PCR using cDNA from *P. berghei* as template and primers P74, P75 (Table S1); followed by cloning into BamHI digested pBCEN5 plasmid using InFusion assembly kit from TaKaRa. The clone was confirmed by PCR, restriction digestion and sequencing. H245A/H247A mutant of PbAMPD (pBCEN5_PbAMPD_H245A/H247A_GFP) was generated by AQUA cloning protocol as described earlier (Beyer et al., 2015). Partial amplicons were generated using primer pairs P76, P79 and P77, P78 (Table S1) and pBCEN5_PbAMPD_GFP vector as template.

### 3.8 Cultivation of *P. falciparum, P. berghei* and transfection in *P. berghei*

Fresh blood (O+ve blood group) was collected from healthy human volunteers and stored in the presence of the anticoagulant, acid citrate dextrose solution at 4 °C overnight. Plasma was separated and discarded. The erythrocytes were washed with 1 × PBS three times and stored at 4°C in incomplete RPMI media containing 25 mM HEPES at 50% hematocrit in the presence of the antibiotic gentamycin 2.5 *μ*g mL*^−^*^1^. *P. falciparum* parasites were grown in complete RPMI medium (20 mM glucose, 0.2% sodium bicarbonate, 0.5% Albumax and 50 *μ*M hypoxanthine) at 5% hematocrit under micro-aerophillic conditions using candle jar setup at 37 °C. Parasitemia was checked regularly by making Giemsa stained smears. 5-10 % parasitemia was maintained and media change was given daily (Trager and Jensen, 1976).

Glycerol stock of wild type *P. berghei ANKA* parasites was injected to a healthy male BALB/c mouse (6-8 weeks old) through intra-peritoneal mode. The parasitemia was monitored by drawing blood from tail snip and observing Giemsa stained blood smears under the microsope (100 × oil immersion objective). Transfection of the parasites was done by following the protocol described by Janse *et al.,* (Janse et al., 2006) using Amaxa 4D nucleofector (P5 solution and FP167 program). Drug resistant parasites were harvested in heparin solution (200 units mL*^−^*^1^) made in incomplete RPMI medium. Glycerol stocks were made by mixing 300 *μ*L of 30 % glycerol and 200 *μ*L of the harvested blood and stored in liquid nitrogen.

### 3.9 Genomic DNA isolation and genotyping of transgenic parasites

The harvested parasites were treated with cold 1 × erythrocyte lysis buffer for 5 minutes at 4 °C and centrifuged at 500 × g for 10 minutes. The supernatant was discarded and the pellet was used for either Western blotting or DNA isolation immediately or stored at −80 °C for future use. For DNA isolation, the parasite pellet was resuspended in 350 *μ*L of TNE (Tris, NaCl and EDTA) buffer. To this 10 *μ*L (10 mg mL*^−^*^1^ stock) RNAseA and 50 *μ*L of 10 % SDS were added and volume was made up to 0.5 mL with TNE buffer. This was incubated at 37 °C for 10 minutes followed by pronase treatment for 1 hour (10 *μ*L of 10 mg mL*^−^*^1^ stock). 1 mL of Tris-saturated phenol was added to the sample and mixed gently by inverting. The sample was centrifuged for 5 minutes at 10000 × g and the supernatant was carefully removed and treated with 1 mL of 1:1 phenol-chloroform solution and centrifuged for 5 minutes at 10000 × g. The resultant supernatant was treated with 1 mL of chloroform-isoamyl alcohol solution and centrifuged at 10000 × g for 5 minutes. The supernatant was mixed with 1/10 volume of 3 M sodium acetate (pH 4.5) and 1 mL of absolute ethanol. Sample was mixed by inverting gently in order to precipitate the DNA. The sample was centrifuged at 10000 × g for 1 minute and the supernatant was discarded. The DNA pellet was washed once with 70 % ethanol solution, air dried and resuspended in 50 *μ*L DNAse free water. This was used for genotyping of parasites by PCR. Southern blotting was done on DNA from knockout parasites by following standard protocols (Sambrook and Russell, 2006). *P. berghei* DHFR 3’ UTR was PCR amplified using primers P71, P72. This PCR product was used to generate the probe for Southern blotting using random primer kit from BRIT and ^32^P labeled dATP.

### 3.10 Localization studies in *Plasmodium*

For *P. berghei*, parasites were harvested in heparin solution and centrifuged at 2100 × g for 5 minutes and the supernatant was discarded. The cells were resuspended in 1 × PBS containing Hoeschst 33342 (10 *μ*g mL*^−^*^1^) and incubated at room temperature for 15 minutes. The cells were centrifuged at 2100 × g for 5 minutes and the supernatant was discarded. The cells were washed once with 1 × PBS and the pellet was resuspended in 70 % glycerol. Cells were mounted on glass slide using cover slips, then sealed and stored at 4 °C. The cells were observed under oil immersion objective (100 ×) of Ziess LSM 510 Meta confocal microscope.

### 3.11 Determination of growth rate of wildtype and ∆*ampd P. berghei*

Glycerol stocks of wildtype and clone C1 of ∆*ampd P. berghei* were injected to two separate mice and parasitemia was monitored by Giemsa stained smears. Blood was harvested in heparin solution once parasitemia reached 0.5-1 % and 10^5^ parasites were injected into 5 fresh mice for both wildtype and ∆*ampd P. berghei*. Parasitemia was monitored by counting parasites in Giemsa stained smears regularly starting from day 1 post-injection and growth rate was determined by plotting percentage parasitemia on y - axis against time (number of days) on x - axis. Mortality rate among mice infected with either wildtype or ∆*ampd P. berghei* parasites, was also determined by plotting percentage survival of mice on y - axis against time (number of days) on x - axis.

### 3.12 Generation of cell lysates for Western blotting

For yeast, 1 mL of culture was mixed with 200 *μ*L of 70 % TCA and kept at −80 °C for 1 hour. The sample was thawed and centrifuged at 10000 × g and the supernatant was discarded. The pellet was washed with 80 % acetone twice and allowed to air dry. The dried pellet was resuspended in 40 *μ*L of 1× SDS-PAGE loading dye, incubated in boiling water bath for 5 minutes, centrifuged to separate the debris and the supernatant was separated on a 12 % SDS-PAGE. Western blotting was performed by semi-dry transfer method on to PVDF membrane.

For *P. berghei*, blood from mice was harvested in heparinised RPMI medium and RBCs were lysed by treating with 1× erythrocyte lysis buffer (155 mM NH_4_Cl, 12 mM NaHCO_3_ and 0.1 mM EDTA) at 4 °C for 5 minutes. The sample was centrifuged and the supernatant was discarded. The parasite pellet was lysed by resuspending it in equal volumes of 2× SDS-PAGE loading dye in the presence of protease inhibitor cocktail. The sample was incubated in boiling water bath for 5 minutes, centrifuged to separate the debris and the supernatant was separated on a 12 % SDS-PAGE. Western blotting was performed by semi-dry transfer method on to PVDF membrane.

## 4 Results

### 4.1 Expression of AMPD in *P. falciparum* and functional characterization by complemetation

PfAMPD gene (PlasmoDB gene ID: PF3D7_1329400) is 3173 bp in length encompassing 7 introns which upon splicing gives rise to a 2082 bp mRNA encoding a 693 amino acid protein. RT-PCR resulted in a 2.1 kb amplicon matching the description in the database (Fig. S1a).

Full-length recombinant PfAMPD with N-terminal (His)_6_-tag when expressed in *E. coli* showed very high levels of degradation and both the full-length and degraded fragments were found to be present in the insoluble fraction of the lysate. Multiple clones of PfAMPD with (His)_6_-tag at N‐ or C-terminal, GFP tag, truncated versions (∆N59, ∆N94 and ∆N178) were made to produce intact soluble protein. None of these variants showed reduced degradation or increased solubility of the protein (Fig. S1b).

AMPD deficient yeast strain grows normally on either YPD medium or minimal medium but shows a distinct growth defect phenotype when grown in the presence of S-adenosyl methionine (SAM) (Fig. S2b). This is due to the accumulation of AMP in the cells which results in the inhibition of *de novo* purine biosynthetic pathway eventually resulting in depletion of GTP pools. This growth defect phenotype can be rescued by expression of a functional AMPD. This principle was utilized to determine the functionality of PfAMPD. Coding sequence of PfAMPD gene was cloned in yeast expression vector under GAL1 promoter and serial dilution and spotting assay was performed to check the functionality of PfAMPD. Rescue of the growth defect phenotype was not seen when compared to control strain expressing ScAMPD (Fig. 1a). Upon performing RT-PCR it was found that full length transcript was not formed in yeast cells expressing PfAMPD. Instead, an aberrantly spliced product was observed (data not shown). Hence the experiment was performed using a codon harmonised (Angov et al., 2008) PfAMPD (hPfAMPD) gene. A clear rescue of growth phenotype was observed in this case showing that PfAMPD is a functional ortholog of ScAMPD (Fig. 1a).

**Figure 1:**
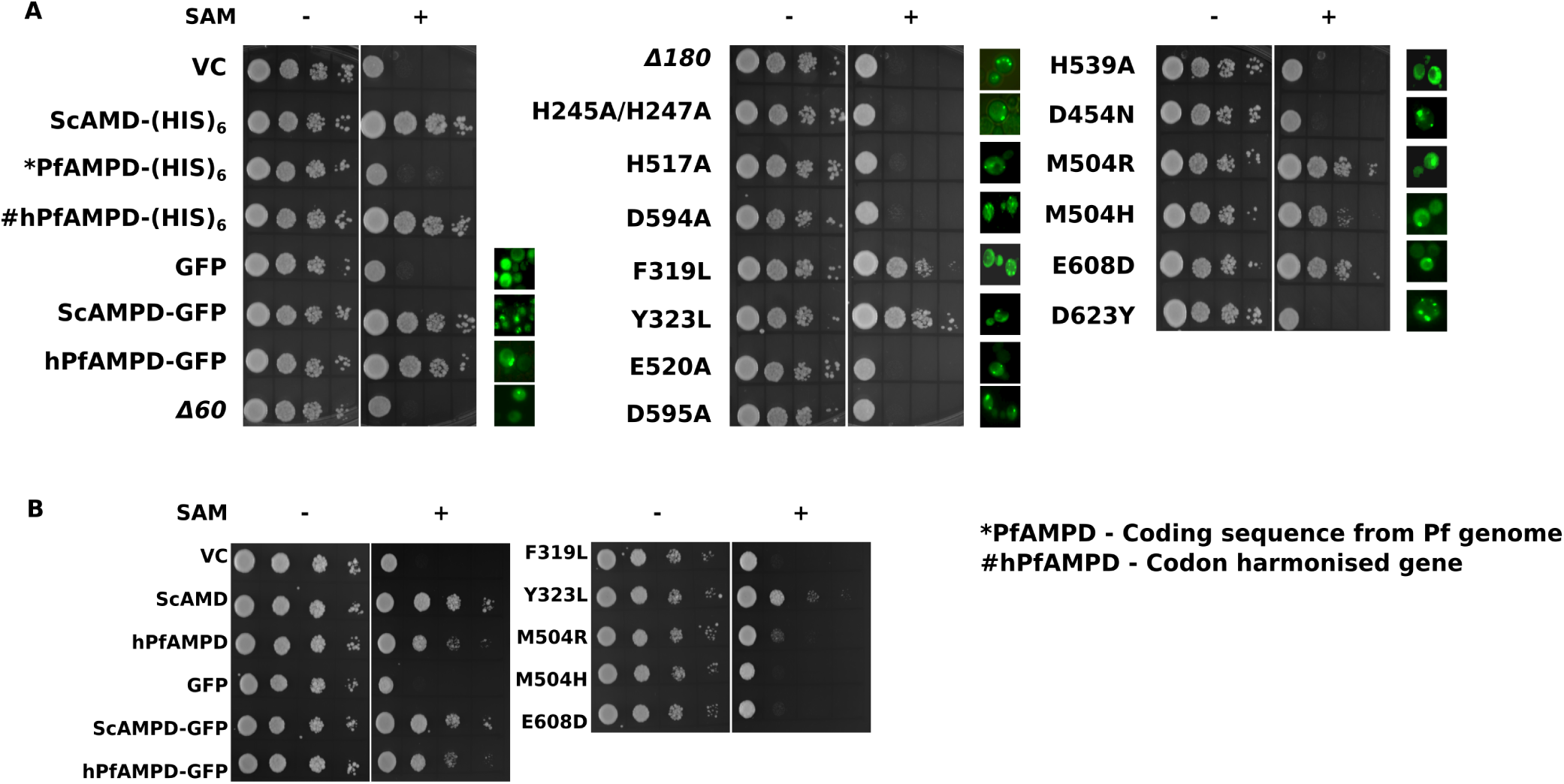
Complementation of AMPD deficiency in *S. cerevisiae*. (A) Ability of truncated and mutant PfAMPDs to rescue the growth defect phenotype of ∆*amd1* yeast strain grown in the presence of SAM was used as a reporter to identify the role of specific residues in enzyme function. Phenotype of mutants was compared to that of vector control (VC - strain containing either pYES2/CT or GFP - pYES2/CT_GFP), ScAMPD, and wildtype PfAMPD expressing cells. The functional complementation assay was performed at 30 °C. Panel A also shows microscopic images of GFP fluorescence as evidence for protein expression. (B) Mutants that were able to rescue the growth defect phenotype at 30 °C were further investigated for their ability to complement at 37 °C and compared with that of wildtype ScAMPD and wildtype PfAMPD expressing cells. (Experiments have been performed in triplicates from two different batches of transformants and image from one experimental replicate is presented).

Although codon harmonised PfAMPD with C-terminal (His)_6_-tag showed functional complementation, full-length protein was not detected by Western blot probed with anti-(His)_6_ antibody. Hence, a C-terminal GFP tagged construct (pYES2/CT_hPfAMPD_GFP) was generated so that it would simultaneously provide proof for the expression of the protein by microscopy and/or by Western blotting probed using anti-GFP antibody. The C-terminal GFP-tagged PfAMPD was also found to functionally complement yeast AMPD deficiency similar to the (His)_6_-tagged protein and moreover, was also detected by Western blot when probed with anti-GFP antibody. Hence, to determine if the GFP tag was conferring stability to the protein, dual tagged PfAMPD constructs (i.e., pYES2/CT_(His)_6__hPfAMPD_GFP and pYES2/CT_hPfAMPD_GFP_(His)_6_) were generated. Western blot was performed using cells expressing various constructs of AMPD i.e., pYES2/CT_hPfAMPD_(His)_6_, pYES2/CT_(His)_6__hPfAMPD_GFP and pYES2/CT_hPfAMPD_GFP_(His)_6_ which were probed with anti-(His)_6_ antibody and pYES2/CT_hPfAMPD_GFP, pYES2/CT_(His)_6__hPfAMPD_GFP and pYES2/CT_hPfAMPD_GFP_(His)_6_ which were probed with anti-GFP antibody. All GFP tagged hPfAMPD constructs were detected by anti-GFP antibody whereas among (His)_6_-tagged constructs pYES2/CT_(His)_6__hPfAMPD_GFP alone was detected by anti-(His)_6_ antibody whereas, the other two constructs (pYES2/CT_hPfAMPD_(His)_6_ and pYES2/CT_hPfAMPD_GFP_(His)_6_) were not detected going to show that the presence of GFP tag at C‐ terminal confers stability to the protein (Fig. S3a). It was also observed that both the constructs (i.e. pYES2/CT_(His)_6__hPfAMPD_GFP and pYES2/CT_hPfAMPD_GFP_(His)_6_) were able to functionally complement yeast AMPD deficiency (Fig. S3b).

### 4.2 Characterization of PfAMPD mutants

AMP deaminases are known to have a divergent N-terminus that is prone to degradation. But, it has been reported that this proteolysis does not affect the activity or regulation of enzyme function (Han et al., 2006; Saint-Marc et al., 2009). A truncated version of AtAMPD has been expressed in insect cell line and the crystal structure solved. This structure showed the presence of a metal centre comprising of Zn^2+^ co-ordinated by H391, H393, H659 and D736 (Fig. S4a) (Han et al., 2006). In addition to this, the report also predicted the role of two hydrophobic residues F463 and Y467 in displacement of the ribose ring (Fig. S4b) and identified E662 and H681 as the putative catalytic residues (Fig. S4c) (Han et al., 2006). However, contribution of these residues to catalysis/ structural stability has not been experimentally validated. By forward genetics approach, residues critical for the functioning of the protein were identified in Arabidopsis (Xu et al., 2005) (D598 makes critical contacts with H659 and the mutant D598N was found to be embryonic lethal) and human (Akizu et al., 2013) (R674H, E778D and D793Y mutants were identified in an exome sequencing study on patients suffering from pontocerebellar hypoplasia) (Fig. S4d). These residues are highly conserved (Fig. 2) and using this information as the basis, site-directed mutagenesis was performed on PfAMPD gene and mutants were characterized by serial dilution and spotting assay (Fig. 1a). It has to be noted that till date, no study using site-directed mutagenesis has been done to validate the role of different residues in AMPD.

**Figure 2:**
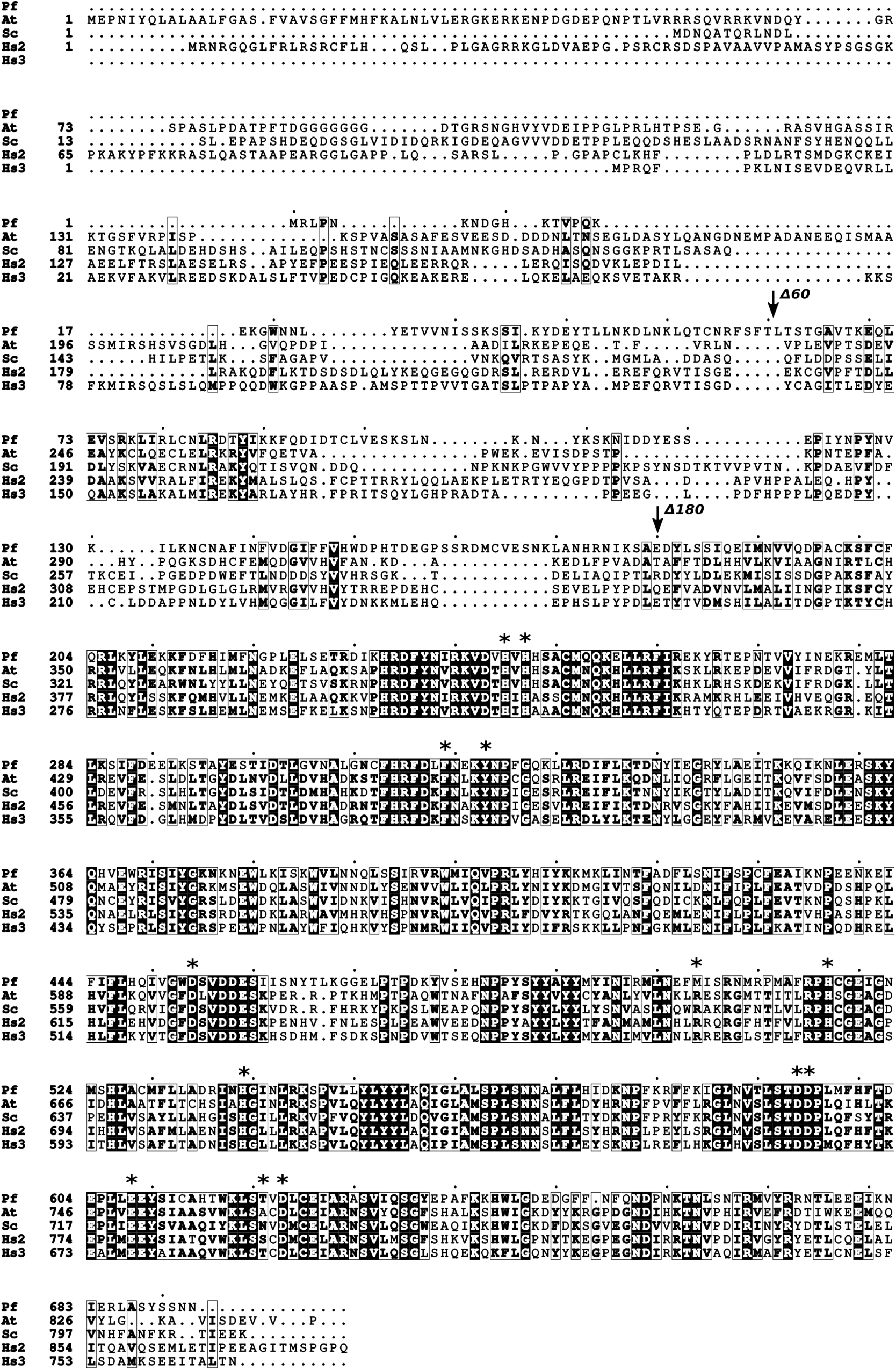
Multiple sequence alignment of AMP deaminase protein sequences. Clustal omega alignment of AMP deaminase sequences from *Plasmodium* (Pf), yeast (Sc), *Arabidopsis* (At) and human (Hs2, Hs3) represented using ESpript 3.0. * indicates residues selected for mutagenesis.

In AtAMPD, deletion of N-terminal 139 residues yielded active enzyme and crystals of diffraction quality. Alignment of AMPD sequences shows that the catalytic core (180 to 680 residues, Pf numbering) is highly conserved (Fig. 2). Both ∆N60 PfAMPD that corresponds to the truncated version of AtAMPD with 139 residues at N-terminus deleted (found to be soluble and used for crystallization), and ∆N180 PfAMPD that corresponds to the catalytic core of the enzyme (Fig. 2) failed to functionally complement yeast AMPD deficiency going to show that in spite of the fact that AMPD sequences have a divergent N-terminus, the N-terminus of PfAMPD might be containing critical residues which are indispensable for activity or structural integrity of the protein. Residues H245, H247, H517 and D594 of PfAMPD that correspond to the residues H391, H393, H659 and D736 of AtAMPD, that co-ordinate with the divalent metal ion at the active site; upon mutation to alanine rendered the protein inactive as cells expressing the mutant proteins failed to grow in the presence of SAM. In AtAMPD structure 2A3L, it was proposed that the residues F463 and Y467 might be involved in displacing the ribose sugar of the ligand coformycin monophosphate (Han et al., 2006). In our study we have found that mutating the corresponding residues in PfAMPD, F319 and Y323 to leucine did not result in complete loss of enzyme activity. However, the F319L mutant did show slightly reduced growth when compared with Y323L or the wild type. In the report by Han *et al.,* (Han et al., 2006) it was proposed that H681 might be the catalytic base and in its absence E662 might perform the same function although both these residues are conserved and are known to be required for catalysis by aminohydrolases in general. Hence, corresponding residues in PfAMPD i.e., H539 and E520 were mutated to alanine and it was found that the cells expressing these mutants failed to grow on SAM indicating that both these residues are indispensable for the activity of the protein and both might be involved in the catalysis. The residue D595 (highlighted in bold and italicized) is present in the conserved motif ‘STD*D*P’ which is seen in all AMP/adenosine deaminases (Han et al., 2006). Mutating this residue to alanine also resulted in loss of activity. A chemical mutagen screen and sequencing study conducted by Xu *et al.,* (Xu et al., 2005) had identified a mutation (D598N) in AtAMPD that resulted in embryonic lethality (Xu et al., 2005). Mutation of the corresponding residue in PfAMPD D454 to alanine resulted in loss of acitivity of the enzyme as observed by lack of rescue of yeast growth defect phenotype. Akizu *et al.,* had identified mutations in human AMPD2 (R674H, E778D and D793Y) which correlated with a neurological disorder called pontocerebellar hypoplasia (Akizu et al., 2013). R674 is conserved in human AMPD2, yeast and *Arabidopsis* AMPD but is a methionine in PfAMPD. This residue was mutated to arginine as well as histidine. Both M504R and M504H mutants were found to be active, but slight growth defect was observed in M504H mutant compared to M504R and wildtype. E608D mutation in PfAMPD corresponding to E778 of human AMPD2 did not have any impact on the protein activity, whereas the D623Y mutant corresponding to D793 of human AMPD2 was inactive.

Expression of GFP-tagged PfAMPD and its mutants at the protein level was determined by Western blotting and also microscopy. Both full-length band corresponding to GFP fusion protein along with a band corresponding to free/degraded GFP was also observed (Fig. S6 a and b). Interestingly, GFP tagged PfAMPD showed multiple localization patterns that included diffused cytosolic, cytoplasmic foci/punctate as well as filamentous pattern (Fig. S6 c-e). The mutants also showed foci/punctate like localization along with the diffused localization pattern (Fig. 1a). Complementation assay was also performed at 37 °C using wild type ((His)_6_-tag and GFP-tag) yeast AMPD and PfAMPD and mutants of PfAMPD which had shown rescue of growth defect phenotype at 30 °C (Fig. 1b). Surprisingly, wild type PfAMPD showed minor yet observable reduction in rescue of growth defect phenotype as compared to the yeast counterpart indicating sensitivity towards increased temperature. Among the mutants, only Y323L mutant of PfAMPD showed rescue of the growth defect phenotype similar to the wildtype, whereas the others completely failed to do so indicating that they might also be playing a role in stabilizing the structure of the protein that is affected at higher temperatures.

### 4.3 Localization studies on AMPD

An episomal construct (pBCEN5_PbAMPD_GFP) where, PbAMPD coding sequence was tagged with C-terminal GFP under PbEF1*α* promoter was used for localization of PbAMPD. When transfected with pBCEN5_PbAMPD_GFP, drug resistant parasites were not obtained. As this result was reproducible with multiple transfection attempts, we hypothesized that increased expression of a stable AMPD over and above endogenous levels in *Plasmodium* might be toxic. Hence, a catalytically inactive mutant (H245A/H247A) of PbAMPD was cloned in pBCEN5 with C-terminal GFP tag and transfected into *P. berghei*. Drug resistant parasites were obtained, genotyped (Fig. 3a) and a diffused cytosolic localization was observed (Fig. 3c). The GFP fusion protein was also detected by Western blotting (Fig. 3b).

**Figure 3:**
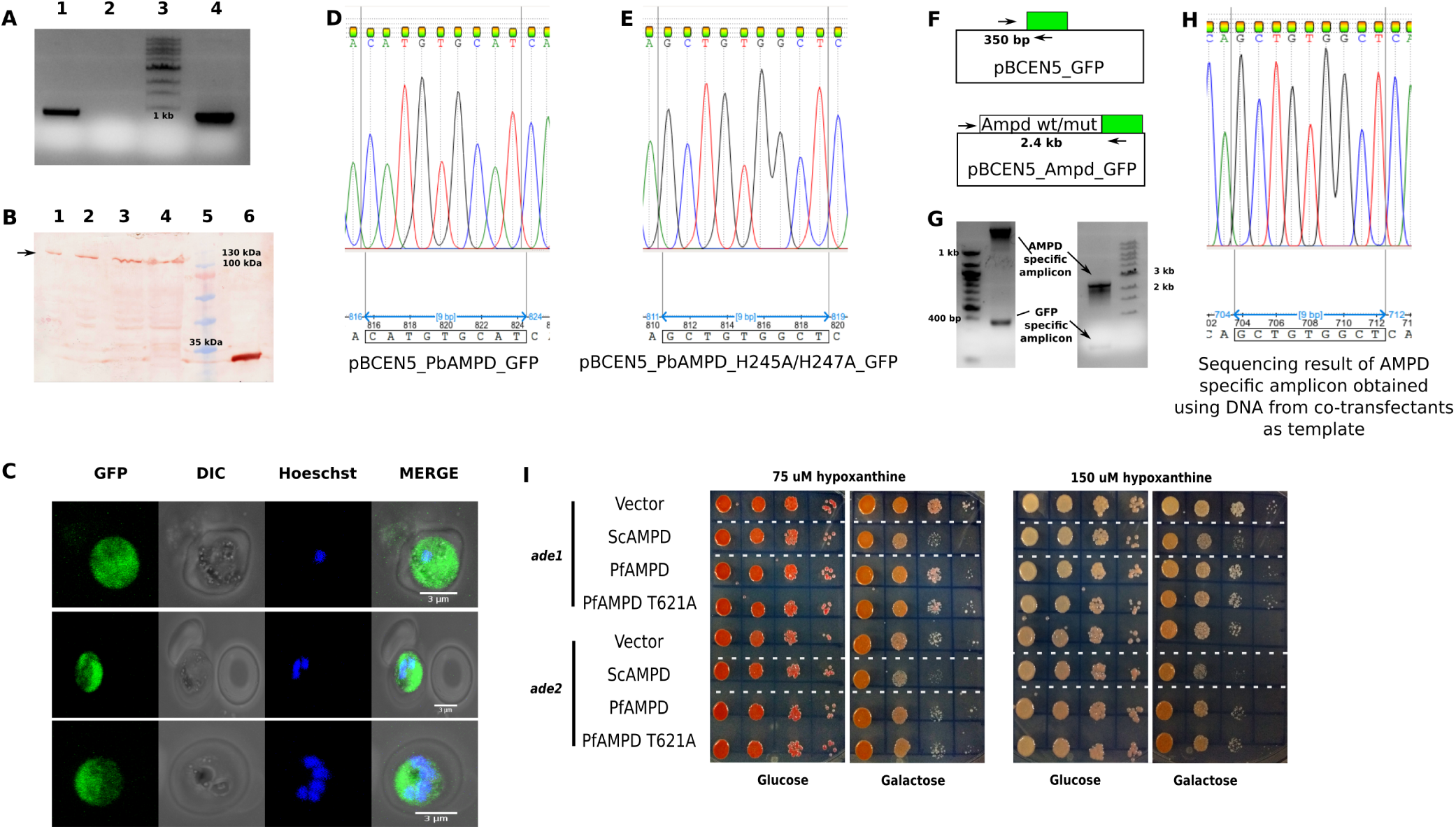
Episomal expression of PbAMPD in *P. berghei*. (A) PCR confirmation of pBCEN5_PbAMPD_H245A/H247A_GFP transfectants. Lane 1, DNA isolated from transfectants as template; lane 2; no template control; lane 3, molecular weight marker; lane 4, +ve control (pBCEN5_PbAMPD_H245A/H247A_GFP). (B) Western blot showing the presence of full length GFP fused PbAMPD_H245A/H247A protein (arrow head) expressed through pBCEN5 in *P. berghei*. Lane 1 to 4, different volumes (5, 10, 20, 40 *μ*L) of lysate from trasfectants expressing PbAMPD_H245A/H247A_GFP fusion protein and lane 6, lysate from *P. berghei* strain expressing GFP alone. (C) Localization of episomally expressed PbAMPD_H245A/H247A_GFP in *P. berghei*. Images were acquired using Ziess LSM 510 META confocal microscope under 100 × oil immersion objective and maximum intensity projections obtained using ImageJ are represented. (D-H) Results pertaining to co-transfection experiments. *P. berghei* was co-transfected with pBCEN5_GFP (vector control), pBCEN5_PbAMPD_GFP, pBCEN5_PbAMPD_H245A/H247A_GFP. Chromatogram D and E show the difference in DNA sequence between the wild type (CAT GTG CAT) and catalytically inactive mutant (GCT GTG GCT). (F) Schematic of the primers selected for genotyping the parasites by PCR. A primer that binds to vector backbone and a GFP internal reverse primer were used that would amplify the DNA segment present in both vector control and PbAMPD (both wildtype and mutant). (G) PCR was performed on DNA isolated from drug resistant parasites which turned out to be positive for both vector control and the gene of interest i.e., PbAMPD (either wildtype or mutant or both). Bands corresponding to GFP alone and PbAMPD (wildtype or mutant or both) are indicated by arrow marks. The left image is of a 1 % agarose gel that was used to visualize the presence lower molecular weight (vector control) PCR product and the right image is that of a 0.8 % agarose gel that was used to visualize the presence of higher molecular weight (PbAMPD gene specific) PCR product. The gene specific PCR product was purified and sequenced to determine whether the sequence corresponds to wildtype PbAMPD or the mutant or a mixed population. As shown in chromatogram H the DNA sequence of transfectants bearing pBCEN5 plasmid containing PbAMPD gene turned out to be the mutant (GCT GTG GCT) and not the wildtype. (I) ∆*ade1* and ∆*ade2* strains were transformed with vectors expressing ScAMPD or PfAMPD or PfAMPD-T621A mutant and grown on minimal medium containing 75 or 150 *μ*M hypoxanthine as purine source. Glucose was used to repress protein expression and galactose for induction. Cells episomally expressing ScAMPD displayed a growth defect phenotype when compared to vector control or PfAMPD (wildtype or mutant) in both strains and at both concentrations of hypoxanthine (highlighted between white dashed lines). Appearance of pink color in the colonies is due to the accumulation of P-ribosylaminoimidazole under purine limiting conditions.

### 4.4 AMPD overexpression in *P. berghei* is toxic

As mentioned above, drug resistant parastites were not obtained when wildtype *P. berghei* was transfected with pBCEN5 expressing PbAMPD-GFP wild type gene in spite of multiple attempts. Whereas, transfection with pBCEN5 expressing PbAMPD_H245A/H247A_GFP mutant gene yielded drug resistant and GFP positive parasites. This suggests that increased levels of AMPD activity might be toxic to the cells and hence the inability to obtain drug resistant parasites. To confirm this inference, pBCEN5_GFP (vector control), pBCEN5_PbAMPD_GFP and pBCEN5_PbAMPD_H245A/H247A_GFP were mixed in equal amounts (2.5 *μ*g each) and co-transfected in to wildtype *P. berghei* which resulted in a mixed population of drug resistant parasites that were PCR positive for both vector control plasmid and plasmid containing either or both PbAMPD wildtype and mutant genes. Sequencing of the AMPD specific PCR product showed the presence of only the plasmid carrying the mutant gene and that the plasmid carrying the wildtype gene was not retained in the drug resistant parasite population (Fig. 3d-h). This confirms that expression of functional PbAMPD above the endogenous levels is toxic to the parasites.

A supporting evidence for the above mentioned inference was also observed in yeast strains lacking functional *de novo* purine biosynthetic pathway. ∆*ade1* and ∆*ade2 yeast* strains are deficient in enzymes of *de novo* purine biosynthetic pathway (N-succinyl-5-aminoimidazole-4-carboxamide ribotide synthetase and phosphoribosylaminoimidazole carboxylase) and hence have been used as mimics of the *Plasmodium* parasite in the context of nucleotide metabolism. Survival of these strains is conditional to the presence of purine precurssors such as hypoxanthine or adenine in the medium which are used to generate AMP. In such a scenario, when an enzyme that catabolizes AMP is over expressed, it results in a futile cycle and compromises the cell viability that will manifest as a growth defect phenotype. Both ∆*ade1* and ∆*ade2* strains grown under conditions with limited quantity of purine, showed growth defect phenotype upon episomal expression of ScAMPD over and above the endogenous levels. Interestingly, episomal expression of PfAMPD in ∆*ade1* and ∆*ade2* strains grown under similar conditions did not result in growth defect phenotype (Fig. 3i). In addition to expression of wildtype PfAMPD protein, T621A mutant (corresponds to a hyperactive mutation observed in human AMPD3 (Hortle et al., 2016)) was also expressed in ∆*ade1* and ∆*ade2* strains but a growth defect phenotype was not observed in this case also.

### 4.5 AMPD is non-essential during asexual stages in *P. bghei*

∆*ampd P. berghei* parasites were generated using linear pJAZZok vector with homology arms flanking the marker gene, generated by recombineering strategy (Godiska et al., 2009; Pfander et al., 2011). The knockout parasites were genotyped by PCR and Southern blotting (Fig. 4), and by limiting dilution three clonal populations (C1, C2 and C3) were obtained that were genotyped by PCR (Fig. S8). One of the clonal populations (C1) was used for assessing the growth phenotype. Comparison of blood-stage growth rate measurements between wildtype and ∆*ampd P. berghei* parasites in mouse showed no significant difference (Fig. 5), thus establishing that the gene is not essential for the parasite during the intra-erythrocytic stages.

**Figure 4:**
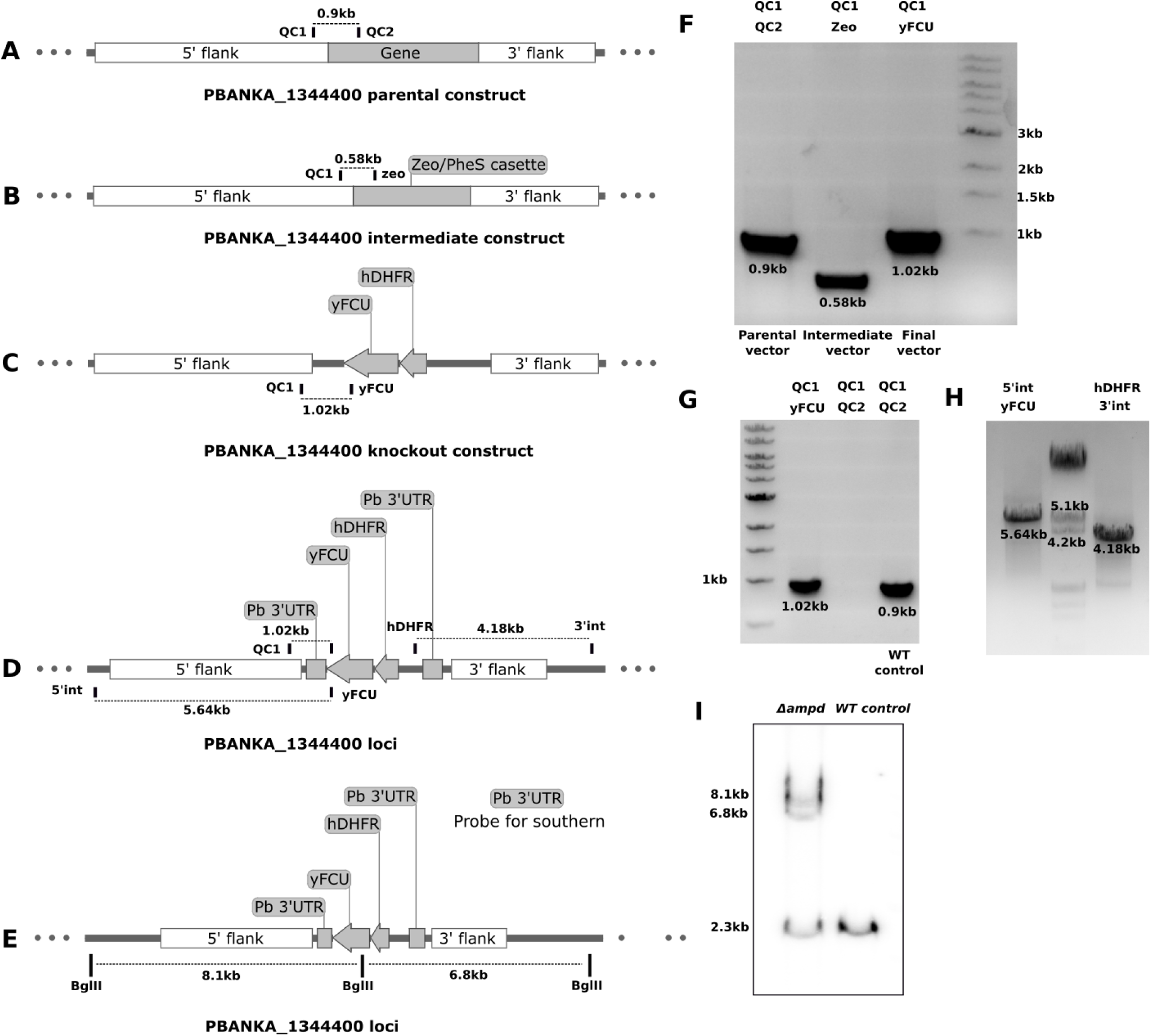
Generation of *P. berghei* ∆*ampd* parasites. (A-D) Schematic representation of PbAMPD parental, intermediate and final knockout constructs and PbAMPD loci after integration. Primers are indicated in the schematic by vertical bars and expected PCR product size is represented by the line between specific primer pairs. (E) Schematic representation of the restriction map of PbAMPD loci with BglII enzyme sites indicated by vertical bars. (F) PCR confirmation of parental, intermediate and final knockout constructs. (G and H) Genotyping of the P. berghei strain for integration of cassette in the correct loci. (I) Genotyping of the strain by Southern blotting using Pb 3’UTR probe.

**Figure 5:**
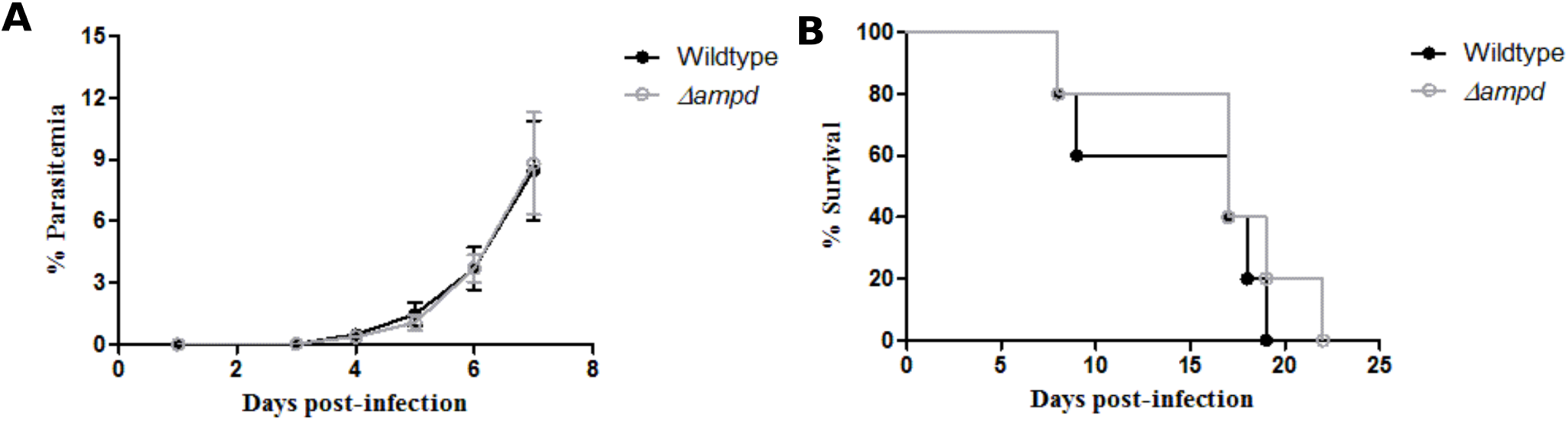
Growth phenotype of wildtype and ∆*ampd P. berghei* parasites. (A) Comparison of growth rates of intra-erythrocytic wildtype and ∆*ampd P. berghei* parasites in mice. The data represents mean *±* s.d. values obtained from five mice each infected with 10^5^ wildtype or ∆*ampd P. berghei* parasites. Statistical analysis was done using Student's unpaired t-test using Graph pad Prism V5. (B) Percentage survival of mice (n=5) infected with either wildtype or ∆*ampd P. berghei* parasites. The survival curves were found to be not significantly different according to Log-rank (Mantel-cox) test (P value = 0.4157) and Gehen-Breslow-Wilcoxon test (P value = 0.5991).

## 5 Discussion

AMPD belongs to amidohydrolase superfamily of enzymes and is a part of purine nucleotide cycle involved in the cyclic interconversion of AMP and IMP. Studies on *Arabidopsis* and human AMPD have lead to the elucidation of the critical role of this enzyme in development, as impairment of its function affects AEC, as well as protein production. To our knowledge, till date, there were no reports on comprehensive characterization of any parasitic protozoan AMP deaminase. Characterization of adenylosuccinate synthetase (ADSS) and adenylosuccinate lyase (ASL) from the malarial parasite *Plasmodium falciparum* have been reported earlier (Jayalakshmi et al., 2002; Eaazhisai et al., 2004; Bulusu et al., 2009; Mehrotra et al., 2010). These two enzymes that constitute one arm of PNC and involved in AMP generation have been found to be essential for the parasite's survival (Sanderson and Rayner, 2017). AMPD constitutes the second arm of PNC, regenerating IMP from AMP and had remained unexplored. This prompted us to conduct investigations towards elucidating the physiological role of AMPD in *Plasmodium*.

Our prime objective of performing biochemical characterization on recombinant PfAMPD was met with numerous hurdles owing to its insoluble and unstable nature. When expressed in *E. coli,* full length as well as truncated versions of PfAMPD showed a characteristic degradation pattern and all the fragments were present as inclusion bodies. Attempts to obtain soluble/stable protein by co-expressing with chaperones as well as refolding strategies were futile. Multiple expression systems and strategies were employed which included yeast (*S. cerevisiae* and *Pichia pastoris* using pYES2/CT and pGAPZ vectors, respectively), and *in vitro* cell-free protein synthesis system. None of these strategies were helpful in obtaining soluble and stable PfAMPD (data not shown). When expression was attempted in *S. cerevisiae* using a codon harmonised gene, the presence of a soluble and functional protein was established based on a growth phenotype readout in a functional complementation assay carried out using ∆*amd1* yeast strain. Although we were able to expresses a functional PfAMPD in a heterologous system, the protein was not detected by Western blot when yeast cell lysates were probed with anti-(His)_6_ antibodies. Hence, to provide evidence for *‘cause (protein expression) and consequence (functional complementation)’* correlation, we generated a PfAMPD-GFP fusion construct. Using this construct, expression of protein was demonstrated both by microscopy and Western blot. The observation that only the GFP-tagged protein can be detected by Western blot and not the (His)_6_-tagged counterpart indicated that the addition of GFP-tag had conferred stability to PfAMPD. In-spite of successfully detecting full length protein on Western blot, we were unable to purify the protein in sufficient quantities for *in vitro* assays and hence had to rely on the use of functional complementation assay for further characterization of the protein.

The crystal structure of AtAMPD (PDB ID 2A3L) was used for addressing structure-function relationships in the *Plasmodium* enzyme. Our study provides a comparative analysis of similarities and differences between yeast, *Arabidopsis*, human and *P. falciparum* AMPD. Establishing the essentiality of the divergent N-terminus and identification of structurally and catalytically important residues was performed using functional complementation assay on truncated protein and site-directed mutants. PfAMPD is different from any other AMPD as the N-terminal truncated version was found to be inactive, where as N-terminal truncated versions of other AMPDs (yeast, *Arabidopsis*, human) have been reported to be active. In comparison with the yeast enzyme, PfAMPD shows a mild temperature sensitive phenotype (when grown in the presence of SAM at 37 °C) which becomes distinct in the mutants F319L, M504R, M504H and E608D indicating the role of these residues in providing structural stability. Residues identified as vital for functioning of human AMPD2 were found to be important for the *P. falciparum* counterpart as well.

Attempts to express GFP-tagged PbAMPD from an episomal vector met with failure as multiple transfection attempts did not yield drug resistant parasites. *Plasmodium* is completely dependent on the host for its purine source and operates predominantly via the HGPRT route for making IMP, which is then converted to AMP via the ADSS and ASL pathway and GMP via the IMPDH and GMPS reactions. AMP thus generated is utilized for various cellular processes including DNA synthesis during cell division. In such a metabolic context, increased expression of an enzyme that catbolizes AMP might result in the futile cycling of AMP/IMP interconversion probably leading to a deficit of AMP that would have deleterious effect on actively dividing cells. This explains our inability to get drug resistant parasites when transfected with vector expressing active AMPD that leads to higher levels of the enzyme in the cell. Whereas, drug resistant parasites were obtained when transfection was done with pBCEN5 expressing catalytically inactive mutant of PbAMPD. In addition to this, when pBCEN5 plasmid expressing either GFP alone or GFP-tagged wildtype PbAMPD or GFP-tagged mutant PbAMPD were co-transfected into *P. berghei*, a mixed population of transfectants that were positive for pBCEN5_GFP and pBCEN5_PbAMPD_H245A/H247A_GFP (inactive mutant) were obtained. This, confirms our inference of cytotoxicity upon AMPD overexpression. Due to non-availability of inducible episomal expression systems in *Plasmodium* we resorted to the yeast model to provide additional evidence for the detrimental nature of increased AMPD expression. We employed ∆*ade1* and ∆*ade2* strains which are very well characterized purine auxotrophs lacking functional *de novo* pathway and solely depend on purine salvage by APRT or adenosine kinase or HGPRT (Fig. S7c). These cells were grown on minimal medium containing only hypoxanthine as purine source, thereby, mimicking the *Plasmodium* parasite in context of purine nucleotide salvage. Under these growth conditions when AMPD was episomally expressed, we observed growth defect when compared to the vector control or cells grown under repressive conditions (Fig. 3i). Although growth defect was seen only in cells episomaly expressing ScAMPD, the lack of this phenotype in cells expressing PfAMPD might reflect on the catalytic efficiencies of the two enzymes and heterologous context of the host expression system. These observations in yeast and in *P. berghei*, in conjunction with a study on a hyperactive mutation in the erythrocyte isoform of AMPD resulting in decreased RBC half life, demonstrates that regulation of this enzyme activity is critical for survival in cells incapable of *de novo* purine biosynthesis. It should be noted that the T621A mutation in PfAMPD which corresponds to the hyperactive mutation reported in the erythrocytic isoform of AMPD, also failed to show growth defect phenotype when episomally expressed in ∆*ade1* and ∆*ade2* strains, although it was found to be active based on the complementation assay in AMPD knockout strain (Fig. S7d and 3i). The critical aspect that determines the cell viability in this scenario would be the relative fluxes between AMP generating arm including HGPRT, ADSS and ASL and the AMP depleting arm of AMP deaminase. From our observation in the yeast model it is evident that the AMPD arm dominates the other arm (HGPRT, ADSS and ASL), upon increased expression of AMPD. Therefore, we conclude that increased expression of AMPD in an organism solely dependent on hypoxanthine salvage like *Plasmodium* might be toxic as it perturbs the AMP homeostasis by depleting AMP levels. Interestingly, episomal expression of ASL in *P. falciparum* and yeast adenosine kinase in *P. berghei*, both of which would increase the flux of AMP production was found to be non-lethal (unpublished data). As AMPD is known to be allosterically modulated by ATP, GTP and phosphate (Merkler et al., 1989; Han et al., 2006), allosteric activators of *Plasmodium* AMPD could exhibit anti-parasitic activity and development of an assay methodology for the identification of such molecules would be useful.

*Plasmodium* has two isoforms of adenylate kinase (AK1 and AK2) and two isoforms of adenylate kinase like protein (ALP1 and ALP2). The loci of both AK and ALP were found to be refractory to knockout in *P. falciparum*, whereas in *P. berghei* AK1 was found to be essential while, AK2 was dispensable during asexual stages (Sanderson and Rayner, 2017). Since AMPD activity can in-turn modulate AK activity, essentiality of this gene was investigated in *P. berghei*. AMPD gene was successfully deleted in *P. berghei,* and the transgenic parasites did not show any difference in phenotype in comparison with wildtype counterpart during intra-erythrocytic stages. Hence, AMPD is not essential for the parasite's survival in asexual stages. It has to be noted that in a recent study by Zhang *et al.,* the gene loci was found to amenable for disruption in *P. falciparum* using piggy-bac transposon mediated saturation mutagenesis (Zhang et al., 2018). The growth defect seen in ∆*amd1* yeast strain manifests only in the presence of adenine or adenosine source (SAM) in external media, which results in the build up of intracellular AMP levels by the action of APRT or adenosine kinase. This can be relieved only by expressing a functional AMPD or by supplementing hypoxanthine or guanine in the media. In mammalian systems, it is the circulating adenosine levels that cause the defective phenotype in ∆*ampd* cells. But for the plasmodial parasite that is harbored inside the RBC, the enzymes APRT and adenosine kinase, which would cause AMP accumulation are absent. Even though the RBC compartment has both APRT and adenosine kinase, and AMP generated via this route can enter the parasite (Cassera et al., 2008); hypoxanthine is the key precursor for purines in *Plasmodium*. An inherently advantageous metabolic wiring seems to have enabled the parasite to survive even in the absence of AMPD (Fig. 6). Nevertheless, the loss-of-function phenotype needs to be examined in other stages of the parasite's life cycle.

**Figure 6:**
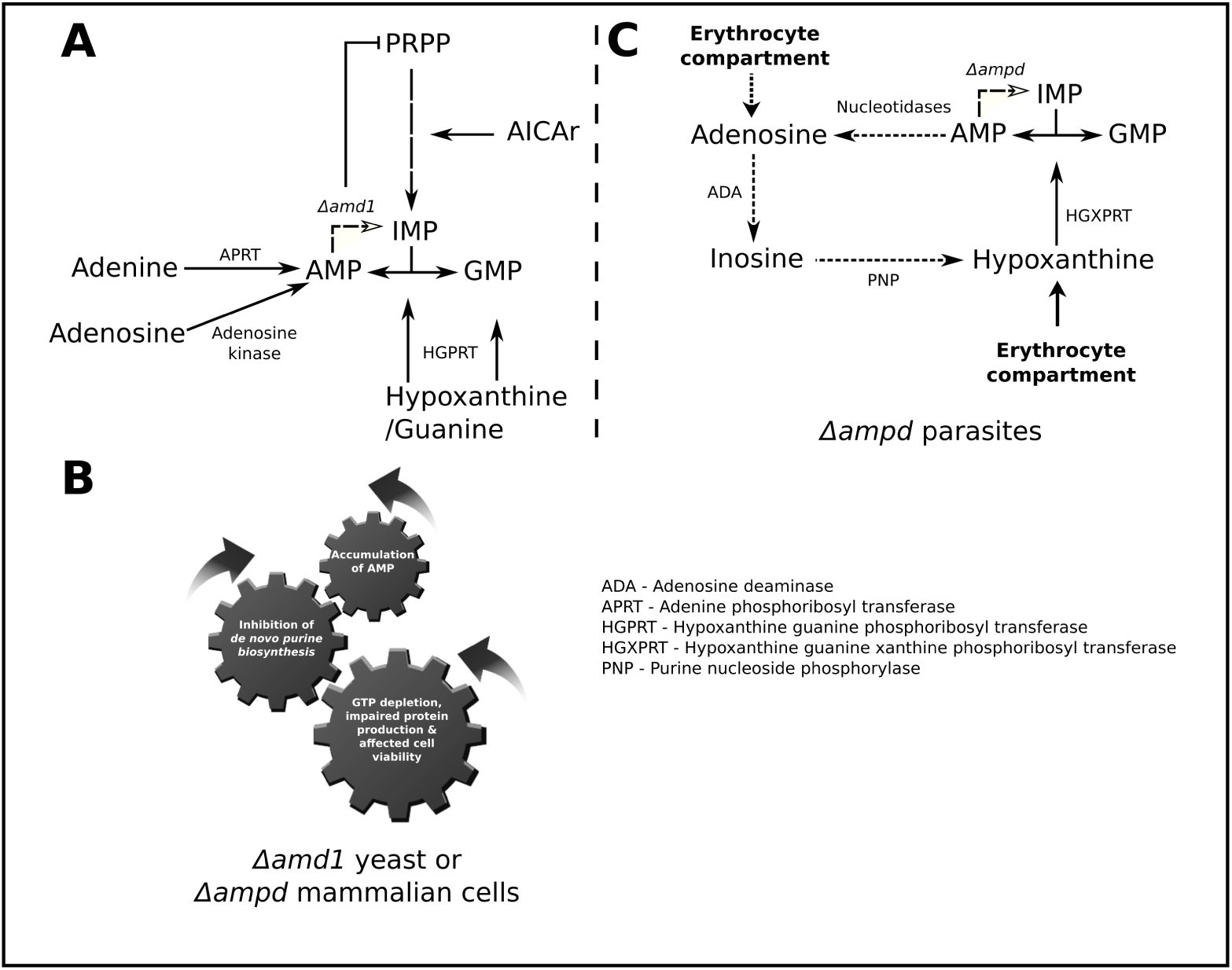
Schematic of AMP metabolism in yeast, mammalian cells and *Plasmodium*. (A) Modes of AMP generation (adenosine phosphoribosyltransferase-APRT and adenosine kinase pathways), consequences of increased AMP concentration in AMPD knockout yeast or mammalian cells and mode of metabolite mediated rescue (HGPRT pathway). (B) The interlinked gears depict how accumulation of AMP eventually manifests as growth defect. (C) Mode of AMP metabolism in ∆*ampd* parasites (note: Enzymes APRT and adenosine kinase which would have lead to AMP accumulation are absent in the parasite). In addition to the salvage of hypoxanthine by HGXPRT other possible route for IMP generation from AMP is also shown (dashed arrow).

In summary, this study is the first report on comprehensive biochemical and physiological characterization of AMPD from an apicomplexan parasite. Apart from providing evidence for the expression of protein in *Plasmodium* by RT-PCR we have also demonstrated that the protein is enzymatically functional by making use of complementation assay in a AMPD knockout yeast strain. In addition, critical residues have been identified that are required for the functionality of this protein and the dispensable nature of this enzyme function in asexual stages of the parasite's life cycle has been established. Having demonstrated that overexpression of AMPD is toxic, one can ascertain that regulation of its activity is vital and it would be interesting to investigate the mode of regulation of this enzyme, employed by the parasite.

## Acknowledgments

This project was funded by; 1) Department of Biotechnology, Ministry of Science and Technology, Government of India. Grant number: BT/PR11294/BRB/10/1291/2014 and BT/PR13760/COE/34/42/2015, 2) Science and Engineering Research Board, Department of Science and Technology, Government of India. Grant number: EMR/2014/001276,COE and, 3) Institutional funding from Jawaharlal Nehru Centre of Advanced Scientific Research, Department of Science and Technology, India. LKN acknowledges CSIR for junior and senior research fellowships.

### Conflict of interest

The authors declare that they have no conflicts of interest with the contents of this article.

### Author contributions

HB and LKN conceived the project and designed the experiments. LKN performed the experiments. LKN and HB wrote the manuscript.

### Ethics statement

Animal experiments involving handling of BALB/c mice were performed by adhering to the standard protocols prescribed by the Committee for the Purpose of Control and Supervision of Experiments on Animals (CPCSEA), a statutory body under the Prevention of Cruelty to Animals Act of 1960 and Breeding and Experimentation Rules of 1998, Constitution of India. The current study (project no. HB002/201/CPCSEA) was approved by Institutional animal ethics committee (IAEC) that comes under the purview of CPCSEA. Whole blood for *P. falciparum* culturing was collected from healthy volunteers with written informed consent.

---

**Table 1:**
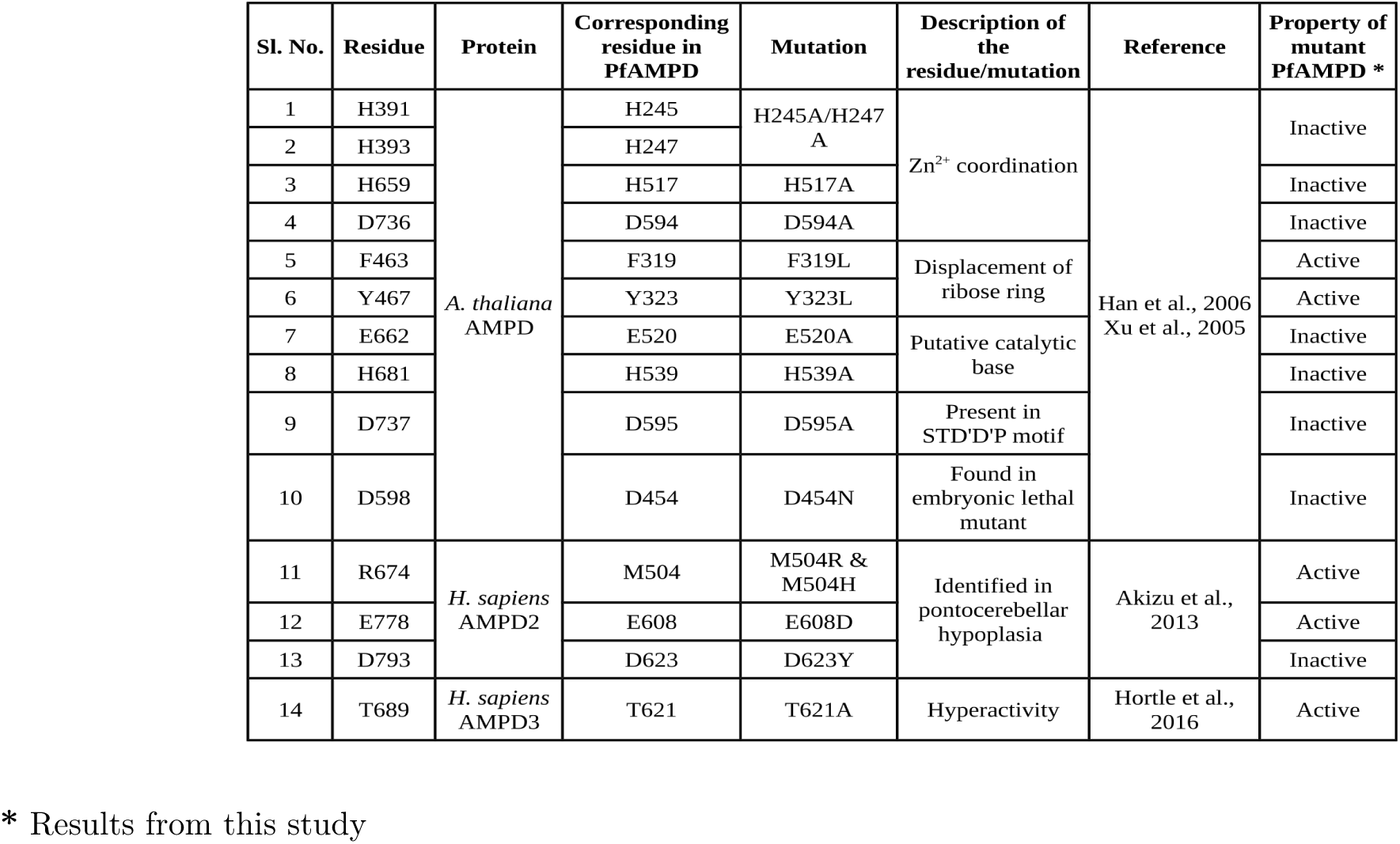
Residues selected for site-directed mutagenesis and their function in enzyme activity

## References

Akizu, N., Cantagrel, V., Schroth, J., Cai, N., Vaux, K., McCloskey, D., et al. (2013). AMPD2 regulates GTP synthesis and is mutated in a potentially treatable neurodegenerative brainstem disorder. Cell 154, 505–517

Angov, E., Hillier, C. J., Kincaid, R. L., and Lyon, J. A. (2008). Heterologous protein expression is enhanced by harmonizing the codon usage frequencies of the target gene with those of the expression host. PloS ONE 3, e2189

Atkinson, D. E. (1968). Energy charge of the adenylate pool as a regulatory parameter. Interaction with feedback modifiers. Biochemistry 7, 4030–4034

Aurrecoechea, C., Brestelli, J., Brunk, B. P., Dommer, J., Fischer, S., Gajria, B., et al. (2008). Plasmodb: a functional genomic database for malaria parasites. Nucleic Acids Res. 37, D539–D543

Bahl, A., Brunk, B., Coppel, R. L., Crabtree, J., Diskin, S. J., Fraunholz, M. J., et al. (2002). Plasmodb: the *Plasmodium* genome resource. An integrated database providing tools for accessing, analyzing and mapping expression and sequence data (both finished and unfinished). Nucleic Acids Res. 30, 87–90

Bahl, A., Brunk, B., Crabtree, J., Fraunholz, M. J., Gajria, B., Grant, G. R., et al. (2003). Plasmodb: the *Plasmodium* genome resource. A database integrating experimental and computational data. Nucleic Acids Res. 31, 212–215

Bazin, R. J., McDonald, G. A., and Phillips, C. (2003). UK Patent GB2373504.

Beyer, H. M., Gonschorek, P., Samodelov, S. L., Meier, M., Weber, W., and Zurbriggen, M. D. (2015). Aqua cloning: a versatile and simple enzyme-free cloning approach. PloS ONE 10, e0137652

Bulusu, V., Srinivasan, B., Bopanna, M. P., and Balaram, H. (2009). Elucidation of the substrate specificity, kinetic and catalytic mechanism of adenylosuccinate lyase from *Plasmodium falciparum*. Biochim. Biophys. Acta 1794, 642–654

Cassera, M. B., Hazleton, K. Z., Riegelhaupt, P. M., Merino, E. F., Luo, M., Akabas, M. H., and Schramm, V. L. (2008). Erythrocytic adenosine monophosphate as an alternative purine source in *Plasmodium falciparum*. J. Biol. Chem. 283, 32889–32899

Chae, S.-C., Fuller, D., and Loomis, W. F. (2002). Altered cell-type proportioning in *Dictyostelium* lacking adenosine monophosphate deaminase. Dev. Biol. 241, 183–194

Chapman, A. G. and Atkinson, D. E. (1973). Stabilization of adenylate energy charge by the adenylate deaminase reaction. J. Biol. Chem. 248, 8309–8312

de Koning-Ward, T. F., Fidock, D. A., Thathy, V., Menard, R., van Spaendonk, R. M., Waters, A. P., et al. (2000). The selectable marker human dihydrofolate reductase enables sequential genetic manipulation of the *Plasmodium berghei* genome. Mol. Biochem. Parasitol. 106, 199–212

Downie, M. J., Kirk, K., and Mamoun, C. B. (2008). Purine salvage pathways in the intraerythrocytic malaria parasite *Plasmodium falciparum*. Eukaryot. cell 7, 1231–1237

Eaazhisai, K., Jayalakshmi, R., Gayathri, P., Anand, R., Sumathy, K., Balaram, H., et al. (2004). Crystal structure of fully ligated adenylosuccinate synthetase from *Plasmodium falciparum*. J. Mol. Biol 335, 1251–1264

Fishbein, W. N., Armbrustmacher, V. W., and Griffin, J. L. (1978). Myoadenylate deaminase deficiency: a new disease of muscle. Science 200, 545–548

Gietz, R. D. and Woods, R. A. (2002). Transformation of yeast by lithium acetate/single-stranded carrier DNA/polyethylene glycol method. Methods Enzymol. 350, 87–96

Godiska, R., Mead, D., Dhodda, V., Wu, C., Hochstein, R., Karsi, A., et al. (2009). Linear plasmid vector for cloning of repetitive or unstable sequences in *Escherichia coli*. Nucleic Acids Res. 38, e88–e88

Han, B. W., Bingman, C. A., Mahnke, D. K., Bannen, R. M., Bednarek, S. Y., Sabina, R. L., et al. (2006). Membrane association, mechanism of action, and structure of *Arabidopsis* embryonic factor 1 (FAC1). J. Biol. Chem. 281, 14939–14947

Hellsten, Y., Richter, E., Kiens, B., and Bangsbo, J. (1999). AMP deamination and purine exchange in human skeletal muscle during and after intense exercise. J. Physiol. 520, 909–920

Hortle, E., Nijagal, B., Bauer, D. C., Jensen, L. M., Ahn, S. B., Cockburn, I. A., et al. (2016). Adenosine monophosphate deaminase 3 activation shortens erythrocyte half-life and provides malaria resistance in mice. Blood 128, 1290–1301

Isackson, P. J., Bujnicki, H., Harding, C. O., and Vladutiu, G. D. (2005). Myoadenylate deaminase deficiency caused by alternative splicing due to a novel intronic mutation in the AMPD1 gene. Mol. Genet. Metab. 86, 250–256

Jahngen, E. G. and Rossomando, E. F. (1986). AMP deaminase in *Dictycostelium discoideum*: Increase in activity following nutrient deprivation induced by starvation or hadacidin. Mol. Cell. Biochem. 71, 71–78

Janse, C. J., Ramesar, J., and Waters, A. P. (2006). High-efficiency transfection and drug selection of genetically transformed blood stages of the rodent malaria parasite *Plasmodium berghei*. Nat. Protocols 1, 346

Jayalakshmi, R., Sumathy, K., and Balaram, H. (2002). Purification and characterization of recombinant Plasmodium falciparum adenylosuccinate synthetase expressed in *Escherichia coli*. Protein Expr. Purif. 25, 65–72

Lanaspa, M. A., Cicerchi, C., Garcia, G., Li, N., Roncal-Jimenez, C. A., Rivard, C. J., et al. (2012). Counteracting roles of AMP deaminase and AMP kinase in the development of fatty liver. PloS ONE 7, e48801

Lushchak, V. I., Smirnova, Y. D., and Storey, K. B. (1998). AMP-deaminase from sea scorpion white muscle: properties and redistribution under hypoxia. Comp. Biochem. Physiol. B, Biochem. Mol. Biol. 119, 611–618

Madrid, D. C., Ting, L.-M., Waller, K. L., Schramm, V. L., and Kim, K. (2008). *Plasmodium falciparum* purine nucleoside phosphorylase is critical for viability of malaria parasites. J. Biol. Chem. 283, 35899–35907.

Mangani, S., Meyer-Klaucke, W., Moir, A. J., Ranieri-Raggi, M., Martini, D., and Raggi, A. (2003). Characterization of the zinc-binding site of the histidine-proline-rich glycoprotein associated with rabbit skeletal muscle AMP deaminase. J. Biol. Chem. 278, 3176–3184

Marotta, R., Parry, B. R., and Shain, D. H. (2009). Divergence of AMP deaminase in the ice worm *Mesenchytraeus solifugus* (annelida, clitellata, enchytraeidae). Int. J. Evol. Biol. 2009

Martini, D., Ranieri-Raggi, M., Sabbatini, A. R., Moir, A. J., Polizzi, E., Mangani, S., et al. (2007). Characterization of the metallocenter of rabbit skeletal muscle AMP deaminase. A new model for substrate interactions at a dinuclear cocatalytic Zn site. Biochim. Biophys. Acta 1774, 1508–1518

Mehrotra, S., Mylarappa, B., Iyengar, P., and Balaram, H. (2010). Studies on active site mutants of *P. falciparum* adenylosuccinate synthetase: insights into enzyme catalysis and activation. Biochim. Biophys. Acta 1804, 1996–2002

Merkler, D. J. and Schramm, V. L. (1990). Catalytic and regulatory site composition of yeast AMP deaminase by comparative binding and rate studies. Resolution of the cooperative mechanism. J. Biol. Chem. 265, 4420–4426

Merkler, D. J. and Schramm, V. L. (1993). Catalytic mechanism of yeast adenosine 5’- monophosphate deaminase. Zinc content, substrate specificity, ph studies, and solvent isotope effects. Biochemistry 32, 5792–5799

Merkler, D. J., Wali, A. S., Taylor, J., and Schramm, V. L. (1989). AMP deaminase from yeast. Role in AMP degradation, large scale purification, and properties of the native and proteolyzed enzyme. J. Biol. Chem. 264, 21422–21430

Meyer, S. L., Kvalnes-Krick, K. L., and Schramm, V. L. (1989). Characterization of AMD, the AMP deaminase gene in yeast. production of amd strain, cloning, nucleotide sequence, and properties of the protein. Biochemistry 28, 8734–8743

Morisaki, T., Gross, M., Morisaki, H., Pongratz, D., Zöllner, N., and Holmes, E. W. (1992). Molecular basis of AMP deaminase deficiency in skeletal muscle. Proc. Natl. Acad. Sci. U.S.A. 89, 6457–6461

Murakami, K. (1979). Amp deaminase from baker's yeast: Kinetic and molecular properties. J. Biochem. 86, 1331–1336

Murakami, K., Nagura, H., and Yoshino, M. (1980). Permeabilization of yeast cells: application to study on the regulation of AMP deaminase activity in situ. Anal. Biochem. 105, 407–413

Ouyang, J., Parakhia, R. A., and Ochs, R. S. (2011). Metformin activates AMP kinase through inhibition of AMP deaminase. J. Biol. Chem. 286, 1–11

Pfander, C., Anar, B., Schwach, F., Otto, T. D., Brochet, M., Volkmann, K., et al. (2011). A scalable pipeline for highly effective genetic modification of a malaria parasite. Nat. Methods 8, 1078

Roy, S., Karmakar, T., Rao, V. S. P., Nagappa, L. K., Balasubramanian, S., and Balaram, H. (2015a). Slow ligand-induced conformational switch increases the catalytic rate in *Plasmodium falciparum* hypoxanthine guanine xanthine phosphoribosyltransferase. Mol. Biosyst. 11, 1410–1424.

Roy, S., Nagappa, L. K., Prahladarao, V. S., and Balaram, H. (2015b). Kinetic mechanism of *Plasmodium falciparum* hypoxanthine-guanine-xanthine phosphoribosyltransferase. Mol. Biochem. Parasitol. 204, 111–120.

Sabina, R. L. and Mahnke-Zizelman, D. K. (2000). Towards an understanding of the functional significance of N-terminal domain divergence in human AMP deaminase isoforms. Pharmacol. Ther. 87, 279–283

Sabina, R. L., Paul, A.-L., Ferl, R. J., Laber, B., and Lindell, S. D. (2007). Adenine nucleotide pool perturbation is a metabolic trigger for AMP deaminase inhibitor-based herbicide toxicity. Plant Physiol. 143, 1752–1760

Saint-Marc, C., Pinson, B., Coulpier, F., Jourdren, L., Lisova, O., and Daignan-Fornier, B. (2009). Phenotypic consequences of purine nucleotide imbalance in *Saccharomyces cerevisiae*. Genetics 183, 529–538

Sambrook, J. and Russell, D. W. (2006). The condensed protocols from molecular cloning: a laboratory manual

Sanderson, T. and Rayner, J. C. (2017). Phenoplasm: a database of disruption phenotypes for malaria parasite genes. Wellcome open research 2

Seibert, C. M. and Raushel, F. M. (2005). Structural and catalytic diversity within the amidohydrolase superfamily. Biochemistry 44, 6383–6391

Shumate, J. B., Katnik, R., Ruiz, M., Kaiser, K., Frieden, C., Brooke, M. H., et al. (1979). Myoadenylate deaminase deficiency. Muscle & Nerve 2, 213–216

Trager, W. and Jensen, J. B. (1976). Human malaria parasites in continuous culture. Science 193, 673–675

Waarde, A. V. and Kesbeke, F. (1981). Regulatory properties of AMP-deaminase from lateral red muscle and dorsal white muscle of goldfish, Carassius auratus (l.). Comp. Biochem. Physiol. B 69, 413–423

WHO (2017). World malaria report.

Xu, J., Zhang, H. Y., Xie, C. H., Xue, H. W., Dijkhuis, P., and Liu, C. M. (2005). Embryonic factor 1 encodes an AMP deaminase and is essential for the zygote to embryo transition in Arabidopsis. Plant J. 42, 743–758

Yoshino, M., Murakami, K., and Tsushima, K. (1979). AMP deaminase from baker's yeast. Purification and some regulatory properties. Biochim. Biophys. Acta 570, 157–166

Zhang, M., Wang, C., Otto, T. D., Oberstaller, J., Liao, X., Adapa, S. R., et al. (2018). Uncovering the essential genes of the human malaria parasite *Plasmodium falciparum* by saturation mutagenesis. Science 360, eaap7847

